# Environmental and spatial drivers shape Odonata diversity in an Amazonian Indigenous Territory

**DOI:** 10.64898/2026.04.13.718303

**Authors:** Jair Costa Miranda-Filho, Joás Silva Brito, Jady Vivia Almeida Santos, Yan Campioni Cavalcante Dantas, Francisco Maciel Barbosa-Santos, Fábio Santos-Silva, Gabriel Martins Cruz, Beatriz Luz-Silva, Paulo Geovani da Silva Gomes, Erival Prata, Raphael Ligeiro, Francieli F. Bomfim, Kiarasy Kaiabi Panara, Korakoko Panara, Sewa Panara, Sakre Panara, Karapow Panara, Kwakore Panara, Sopoa Panara, Nhasykiati Panara, Pâssua Pri Panara, Pente Panara, Tepakriti Panara, Antonio Ramyllys Oliveira Costa, Lais Sarlo, Bruno Coutinho, Renata Pinheiro, Paulo Junqueira, Isabella Millena Alves Evangelista, Luciano Fogaça de Assis Montag, Thaisa Sala Michelan, Leandro Juen

**Author notes:** Corresponding Author: Jair da Costa Miranda Filho.

## Abstract

Amazon streams are increasingly threatened by land-use change, yet Indigenous Territories represent some of the most effective areas for maintaining habitat integrity and ecological processes in these systems. Understanding how local environmental conditions, landscape context, and spatial structure interact to shape biodiversity within these territories is essential for advancing conservation strategies. Here, we evaluated the relative influence of local habitat, landscape, and spatial predictors on Odonata diversity and identified species-specific ecological thresholds within an Indigenous Territory in the southern Brazilian Amazon. Adult Odonata were sampled in 31 first- to third-order forested streams in the Panará Indigenous Territory, Xingu River basin. Local habitat variables were the main drivers of Odonata community structure, indicating that local habitat integrity and physical stream characteristics strongly influence assemblage composition. In contrast, Zygoptera suborder were primarily structured by spatial predictors, suggesting stronger dispersal limitations and fine-scale spatial processes. Anisoptera suborder showed no significant community-level associations with the predictors, reflecting their broader ecological tolerance and higher dispersal capacity. Our results demonstrate that even within highly conserved Indigenous Territories, subtle environmental gradients and spatial structure shape Odonata assemblages and define ecological thresholds. By integrating community-level and species-specific approaches, this study provides robust evidence of the role of Indigenous lands in sustaining freshwater biodiversity and highlights the value of Odonata as indicators for monitoring ecological integrity in Amazonian streams.

## Introduction

The Amazon basin presents one of the richest biodiversity on Earth, encompassing terrestrial and freshwater fauna and flora (Guayasamin et al. 2024). Amazon freshwater ecosystems play a central role in sustaining regional biodiversity and ecosystem functioning, particularly in forested landscapes where small streams dominate the drainage network (Ferreira et al. 2023). The main drivers of Amazonian freshwater biodiversity include local conditions (e.g. vegetation cover and habitat structure provided by macrophytes) (Thomaz and Cunha 2010; Brasil et al. 2017; Maués-Silva et al. 2024), as well as dispersal-related processes operating across spatial scales. Additionally, amazonian freshwater systems impose important constraints on species’ geographical distributions, limiting dispersal and population establishment (Wallace 1854; Ribas et al. 2012; Guayasamin et al. 2024). Human activities have altered how these drivers affect stream species, changing biodiversity patterns.

In the Brazilian Amazon, human land-use activities (e.g., logging and mining) have increasingly altered freshwater ecosystems, primarily through deforestation and the conversion of native vegetation into agricultural and pasture lands (Faria et al. 2024; Cruz et al. 2025). These changes directly affect key environmental features that structure aquatic communities, particularly in small streams, which are strongly influenced by surrounding landscape conditions (Carvalho et al. 2018; Cruz et al. 2025; 2026). The removal of riparian vegetation, for example, reduces canopy cover, alters light availability, increases water temperature and sediment input, and disrupts the supply of allochthonous resources to the stream channel, ultimately affecting habitat integrity (Dosskey et al. 2010; Juen et al. 2016; Dala-Corte et al. 2020). Additionally, landscape modifications can reduce macrophyte functional diversity, thereby reducing the structural complexity these plants provide and contributing to a decline in associated aquatic biodiversity (Thomaz and Cunha 2010; Bomfim et al. 2023, 2025).

At broader spatial scales, landscape modification reduces environmental heterogeneity and connectivity, potentially influencing the distribution of aquatic organisms (Carvalho et al. 2018; Faria et al. 2024). Given the cumulative effects of local and landscape-scale disturbances on stream ecosystems, identifying mechanisms that maintain the freshwater biodiversity has become a central challenge in Amazonia conservation (Bastos et al. 2021).

In this context, protected areas, including Indigenous Territories, play a pivotal role in biodiversity conservation in the Amazon (Mulongoy and Chape 2004; Azevedo-Santos et al. 2019). Indigenous Territories help maintain natural habitats and ecological processes, supporting the long-term protection of Amazonian ecosystems (Câmara-Leret et al. 2019; Mattos et al. 2024; Athayde et al. 2025). These areas in the Amazon are often characterized by high forest vegetation cover, preserved riparian zones and landscape connectivity. This dynamic is particularly relevant given the importance of aquatic ecosystems for the subsistence of Amazonian Indigenous peoples (Pokhrel et al. 2015; Osborne et al. 2024; Athayde et al. 2025; Façanha et al. 2026).

Despite their recognized importance, empirical assessments of freshwater ecological patterns within Indigenous Territories remain limited (Guerrero-Moreno et al. 2025). There is a persistent gap regarding the insufficient recognition of Indigenous knowledge systems and the underrepresentation of Indigenous Peoples as active agents in biodiversity research, particularly in aquatic ecosystems (Silva-Santos et al., unpublished data; Levis et al. 2024). In this context, the use of reliable bioindicators is essential for better understanding ecological patterns and processes in these environments.

Dragonflies (order Odonata) are widely recognized as effective bioindicators of stream ecosystem condition (Calvão et al. 2016; Brito et al. 2023). Odonata comprises two suborders, Zygoptera and Anisoptera, which are highly mobile predatory insects inhabiting streams, lakes, and ponds. The order currently includes approximately 6,300 described species (Schorr and Paulson 2019) and has high potential for ecological and biomonitoring studies (Miguel et al. 2017; Mendoza-Penagos et al. 2020; Bybee et al. 2021; Córdoba-Aguilar et al. 2023; Goodman et al. 2026). Zygoptera species, which exhibit smaller body size and lower dispersal capacity, are generally more sensitive to canopy removal and habitat degradation. On the other hand, Anisoptera species, although also dependent on structurally complex environments during part of their life cycle, tend to have greater dispersal ability and tolerance to environmental variation (Remsburg et al. 2008; Oliveira-Junior and Juen 2019; Deacon et al. 2024).

Beyond local environmental conditions, the distribution of Odonata species in the Brazilian Amazon is also strongly influenced by biogeographical factors, such as large rivers which act as barriers and contribute to the formation of centers of endemism, limiting species dispersal across the landscape (Juen and De Marco 2012; Brasil et al. 2018; Brito et al. 2024b). Therefore, understanding the relative importance of environmental and spatial drivers is essential for elucidating patterns of Odonata diversity and assessing the role of Indigenous Territories in conserving freshwater biodiversity (Garnett et al. 2018; Amin et al. 2019).

Despite their recognized importance, empirical assessments of freshwater ecological patterns within Indigenous Territories remain scarce, particularly those integrating environmental, spatial, and species-specific responses. Therefore, this study aimed to a) assess how environmental, landscape, and spatial predictors shape patterns of Odonata diversity within an Indigenous territory, b) elucidate species-specific ecological thresholds, and c) provide a comprehensive assessment of species conservation, characterize the conservation status and distribution of recorded species based on publicly available databases (IUCN, ICMBio, and SALVE). Given the contrasting ecological requirements of Odonata suborders and the natural preserved status of streams within Indigenous territory, we hypothesized that: (I) Anisoptera species composition and ecological thresholds would be more strongly associated with less preserved conditions, such as lower forest vegetation cover and greater stream width and depth; on the other hand, (II) Zygoptera, composed mainly of smaller-bodied species, would be associated with higher habitat integrity and features of pristine small streams (e.g., lower width and depth and higher forest vegetation cover); and (III) spatial predictors would exert a stronger influence on Zygoptera due to its ecological and physiological constraints. To test these hypotheses, we combined community-level multivariate analyses with species-specific threshold detection, integrating broad ecological patterns with fine-scale responses along environmental gradients, thereby advancing knowledge of the use of odonates in the biomonitoring of Amazon streams.

## Materials and methods

### Study area

Sampling was carried out in 31 first to third-order streams of the Xingu River basin, located in the Panará Indigenous Territory, which covers approximately 5,000 km² (Fig. 1). The study area lies between the municipalities of Altamira, Pará, and Guarantã do Norte, Mato Grosso, Brazil. The region reaches altitudes of up to 500 m and is characterized by dense ombrophilous forest where the forest canopy reaches approximately 35 m in height and savanna–seasonal forest ecotone vegetation, under a humid tropical climate (*Aw*) according to the Köppen–Geiger classification (Schwartzman and Zimmerman 2005; Peel et al. 2007; Schwartzman 2010). Mean annual precipitation in the region ranges from 1,500 to 2,000 mm, with a pronounced dry season and strong seasonality.

**Fig. 1.**
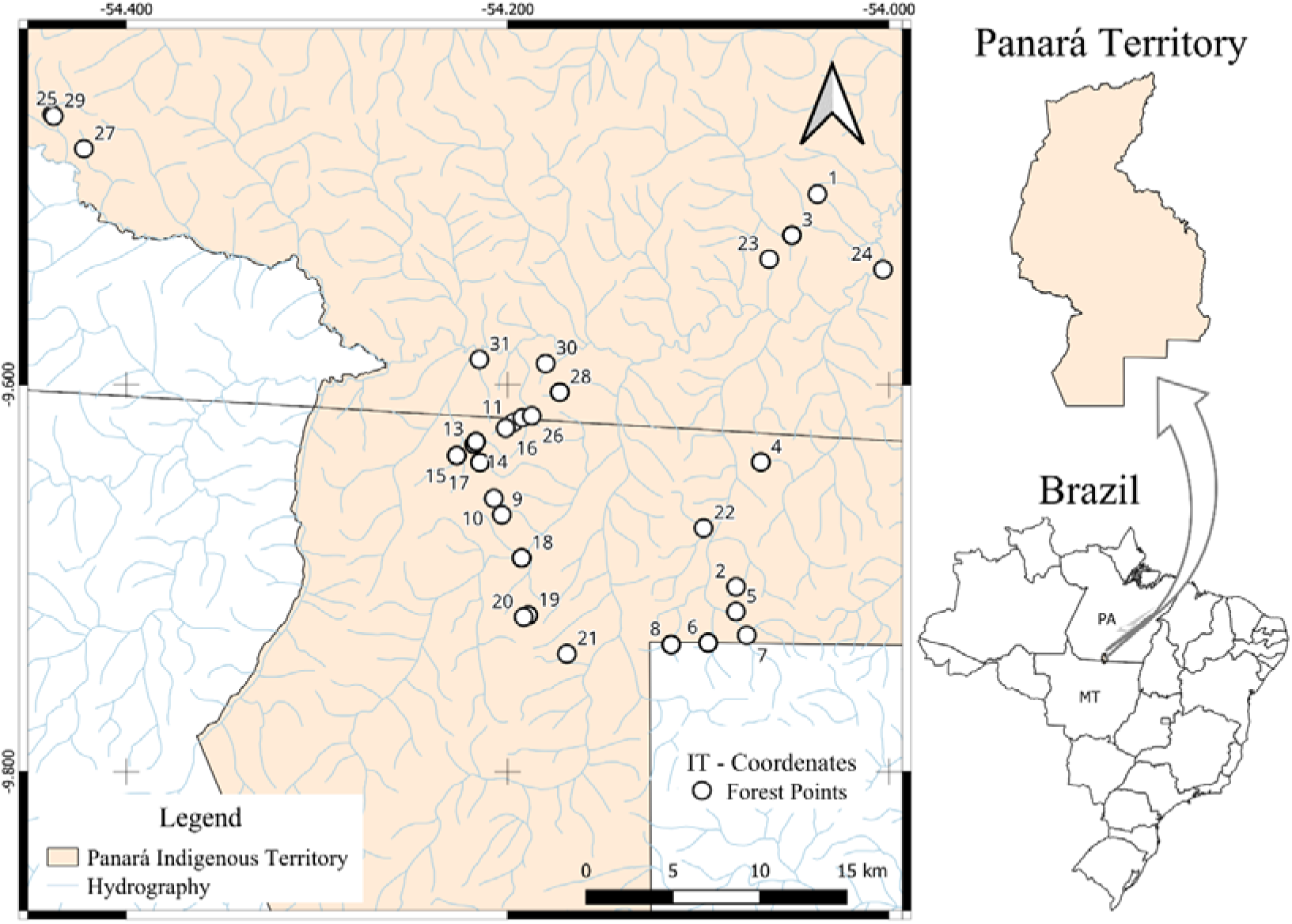
Location of the sampling sites within the Panará Indigenous Territory, between the states of Pará and Mato Grosso, Brazil.

The Panará Indigenous Territory comprises seven villages: *Nâsepotiti*, *Kressã*, *Sankue*, *Sokransan*, *Nãnpôorõ*, *Kotiko*, and *Kanaã* and has an indigenous population of approximately 850 inhabitants (Ewart 2013; ISA 2025). The territory is bordered by three other protected areas, including the *Menkragnoti* and *Terena Gleba Iriri* Indigenous Territory and the Serra do Cachimbo Biological Reserve, as well as by agricultural frontiers dominated by soybean and maize cultivation (Rocha 2015). Traditional land-use practices prevail in the area, including subsistence agriculture and sustainable resource management (Schwartzman 2010).

### Participatory data collection by Panará researchers

This study was conducted in collaboration with the Panará Indigenous People, whose participation served as a central methodological and conceptual component of the research. Eleven Indigenous researchers were actively involved throughout all stages of the project, including the discussion and refinement of the study design, selection of sampling sites, field data collection, specimen triage, and the sharing of knowledge related to the biology, behaviour, and habitat use of Odonata species. This collaborative approach ensured ecologically meaningful and culturally appropriate sampling strategies, while also promoting the co-production of knowledge between Indigenous ecological knowledge and academic science. Such co-production of knowledge strengthens data quality, enhances contextual interpretation of ecological patterns, and contributes to more inclusive and ethical biodiversity research within Indigenous Territories (Fernández-Llamazares et al. 2021; Levis et al. 2018).

### Biological sampling

Adult odonates were sampled using standardized ecological survey protocols (Juen et al. 2025) during the regional dry season (April and May 2025), between 10:00 and 14:00 h, considering the ecophysiological characteristics of these groups (Oliveira-Junior and Juen 2019; Batista et al. 2021). Sampling was performed along 150 m reaches at each stream site using an entomological hand net (40 cm in diameter and 65 cm in depth) (Cezário et al. 2020). Each site was surveyed for 60 minutes under favorable weather conditions to maximize detectability and capture efficiency. This standardized protocol has been widely applied and validated in ecological studies of Amazonian Odonata communities, ensuring comparability among sampling units (Brasil et al. 2017; Oliveira-Junior and Juen 2019; Batista et al. 2021).

After the collection, specimens were stored in paper envelopes and immersed into a 90% alcohol solution. In the laboratory, the specimens were prepared according to Lencioni protocol (2005, 2006), identified using specific taxonomic keys (Lencioni 2005, 2006; Garrison et al. 2006, 2010 for families and genus; and Ris 1930, Borror 1931, 1942; Garrison 1990, Needham et al. 2000; von Ellenrieder 2012; Garrison and von Ellenrieder 2018, for species). When necessary, the collected specimens were sent to specialists to solve identification issues. Finally, after the identification procedures, the specimens were deposited in the Zoology Museum Collection at the Federal University of Pará. To assess the conservation status of the Odonata species recorded in this study, information was compiled by the International Union for Conservation of Nature. (IUCN 2026) Red List and from the Brazilian Red List of Threatened Species. (ICMBio 2026). Additionally, data on species occurrences within Brazilian territory were obtained from the SALVE platform (ICMBio 2026).

### Environmental measurements, land uses and spatial filters

We measured three categories of environmental variables. Individual metric choices were based on the literature of the studies that evaluated the distribution of the order in the biome (Brasil et al. 2018; Resende et al. 2018; Brasil et al. 2020; Brito et al. 2024a):

**1) Local habitat**: aquatic macrophyte species richness (n), depth (cm), and width (m) of the stream channels, and local canopy cover (%). Aquatic macrophyte species richness data were obtained by recording all species present along a 150 m transect. Macrophytes were identified to the lowest possible taxonomic level using specialized literature (Pott and Pott 2000; Fares 2020). Macrophyte species richness was used as a proxy for habitat structural complexity, as multispecies assemblages tend to support greater environmental heterogeneity than monospecific stands (Thomaz and Cunha 2010; Sousa et al. 2025). Water depth and channel width were measured by using a metric PVC pipe (150 cm). Canopy cover was measured by using a convex densiometer, taking four readings in different directions (upstream, downstream, left, and right) (Juen et al. 2016). Additionally, we applied the Habitat Integrity Index (HII), which measures the completeness of the riparian vegetation surrounding streams’ channel and retention devices (e.g., gravel, rocks, wood debris, banks’ leaves) (Nessimian et al. 2008; Brasil et al. 2020a). It varies from 0 (streams degraded or altered) to 1 (preserved conditions) (Nessimian et al. 2008; Brasil et al. 2020b).
**2) Landscape variables**: Landscape proxies, such as forest network, were quantified using land-use and land-cover data obtained from *Projeto MapBiomas*, *Collection 10* (Projeto MapBiomas 2025; https://brasil.mapbiomas.org/), which provides a 10-m spatial resolution product corresponding to the sampling year of each stream. First, we created 30-m-wide linear buffers along the entire upstream dendritic drainage network, which corresponds to the minimum riparian protection width defined by the Brazilian legislation (*Novo Código Florestal, Art. 4°*; https://legis.senado.gov.br/norma/546624/publicacao/15635836). We also delineated the catchment area for each stream. Catchment delineation was based on the geographic coordinates obtained at the most downstream point of each stream site and a digital elevation model (DEM) (INPE; http://www.webmapit.com.br/inpe/topodata/) (Cruz et al. 2026). Finally, we overlaid the linear buffers and catchment areas with the classified land-cover raster to calculate the proportion of forest cover within the network and catchment scales for each stream, respectively. As a result, we had different land uses (forest, grasslands, mosaic, agriculture, and livestock). Given that some of them occurred at only a few sampling sites (fewer than 3), we summarized land uses into two classes: forest and agriculture use. All geoprocessing procedures were performed in *QGIS* (version 3.34), using the *r.watershed* and *r.outlet* algorithms. Additional methodological details are provided in Cruz et al. (2025, 2026).
**3) Spatial variables**: the spatial predictor dataset was generated using spatial eigenfunction analyses (Borcard et al. 2004), based on the Principal Coordinates of Neighbour Matrices (PCNM) (Dray et al. 2006). The PCNM method is based on the geographical distance matrix from the sampling sites (latitude and longitude). We used the minimum spanning tree criterion as the threshold for truncating the geographic distance matrix. At the end, spatial eigenvectors are the main result (PCNMs), whose ecological interpretation is the first PCNMs represent broad-scale predictors, and while the last PCNMs represent fine-scale predictors.

The general information about local habitat and landscape variables is available in Table 1.

**Table 1.** Overall characteristics of the local habitat and landscape measured in the streams sampled inside Panará indigenous lands. HII = Habitat Integrity Index.

| Variables | Mean | Max | Min | SD |
| --- | --- | --- | --- | --- |
| Macrophyte richness | 9.387 | 19.000 | 2.000 | 4.862 |
| Water depth | 18.76 | 48.46 | 6.21 | 9.842 |
| Channel width | 2.409 | 4.080 | 0.96 | 0.756 |
| HII | 0.795 | 0.979 | 0.301 | 0.141 |
| Canopy cover | 90.501 | 100.000 | 5.511 | 19.163 |
| Forest | 9.021 | 94.489 | 0 | 19.215 |
| Agriculture | 9.387 | 19.000 | 2.000 | 4.862 |

### Analytical procedures

Each sampled stream site was treated as an independent sampling unit, totaling 31 samples. Biodiversity patterns were assessed both overall and separately for the suborders Anisoptera and Zygoptera. Additionally, to evaluate the effects of each predictor set while controlling for the others, we performed partial redundancy analysis (pRDA; Borcard et al. 2018), a constrained ordination method analogous to multiple linear regression that assesses the influence of a predictor matrix (X) on a response matrix (Y) while controlling for a third matrix (W). The relative contributions of local habitat, landscape, and spatial predictors to Odonata species composition were quantified with adjusted R² values (Peres-Neto et al. 2006; Borcard et al. 2018).

Prior to the analysis, forward selection was applied to standardized local and landscape variables (forest and agriculture gradients) and to spatial filters (PCNMs) using the forward.sel.par function from the adespatial package (Blanchet et al. 2008). This procedure reduced model complexity by retaining only variables that significantly contributed to explained variation (adjusted R², p < 0.05). Selected variables were subsequently included in the pRDA models. When forward selection retained predictors from a single dataset, standard redundancy analysis (RDA) was applied, with species composition as the response matrix. To generate the triplots depicting the overall associations between Odonata and the predictor variables (environment, landscape, and space), we run global models to create the visuals.

To assess species-specific ecological thresholds and their associations with environmental gradients, we employed Threshold Indicator Taxa Analysis (TITAN; Baker and King 2010). TITAN evaluates changes in species distributions along continuous environmental gradients and identifies thresholds corresponding to shifts in species frequency and relative abundance (Baker and King 2010; Baker et al. 2023). The approach combines two analytical frameworks: (i) Indicator Value analysis (IndVal; Dufrêne and Legendre 1997), which quantifies taxon fidelity (relative frequency) and specificity (relative abundance) across gradient categories; and (ii) Change-Point Analysis (nCPA; King and Richardson 2003), which detects community boundaries along environmental gradients by estimating change points and confidence intervals. Species responses are expressed as standardized z-scores and classified as Z− (negative responses) or Z+ (positive responses) relative to the environmental gradient (Baker and King 2010; Baker et al. 2023).

To perform TITAN, we followed standard analytical requirements: (i) each species had to be represented by at least five individuals, and (ii) these individuals had to occur across at least five sampling sites (Baker et al. 2023). Low frequencies and abundances can compromise TITAN results, leading to less robust estimates. We applied a bootstrapping procedure (n = 100) and random permutations (n = 1000) to assess the robustness of species responses. Species were classified as tolerant or sensitive when both purity and reliability were ≥ 0.90.

All analyses were conducted in the R computational environment (R Core Team 2025) using the RStudio interface. Ecological thresholds were estimated with the *titan* function implemented in the *TITAN2* package (Baker et al. 2023).

## Results

### Assemblage overview

Approximately 800 adult Odonata specimens were collected, representing 49 species distributed across the two suborders: 29 Anisoptera and 20 Zygoptera. In total, 9 families were recorded Coenagrionidae (n = 267), Calopterygidae (n = 227), and Libellulidae (n = 215) being the most abundant. Within Coenagrionidae, *Argia oculata* (n = 98), *Argia collata* (n = 88), and *Acanthagrion gracile* (n = 39) were the most abundant species. In Calopterygidae, *Hetaerina moribunda* (n = 150) and *Hetaerina caja* (n = 13) accounted for the highest numbers of individuals. For Libellulidae, *Erythrodiplax fusca* (n = 39), *Perithemis thais* (n = 25), and *Orthemis discolor* (n = 13) were the most prevalent species (Table 2).

**Table 2.**
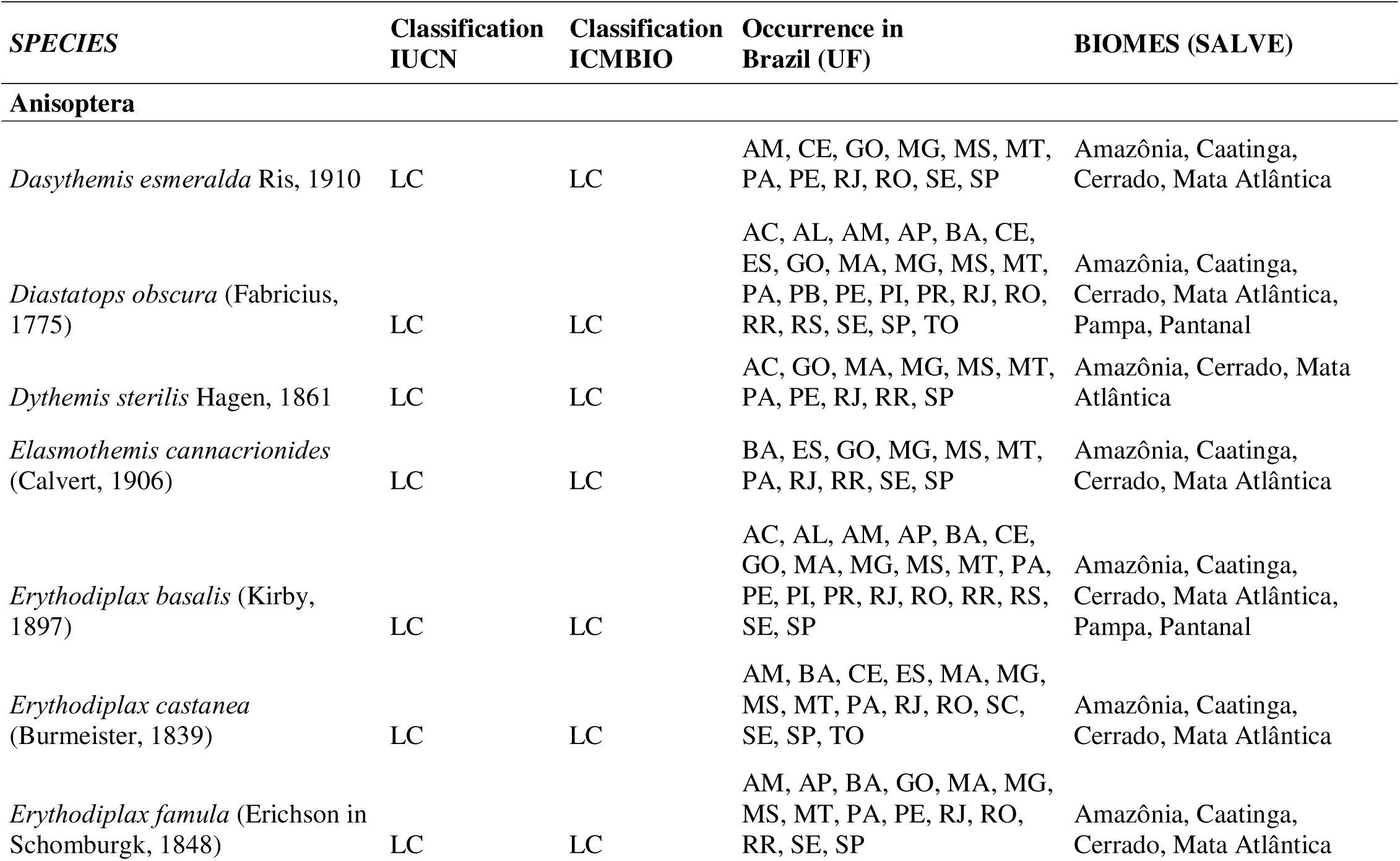
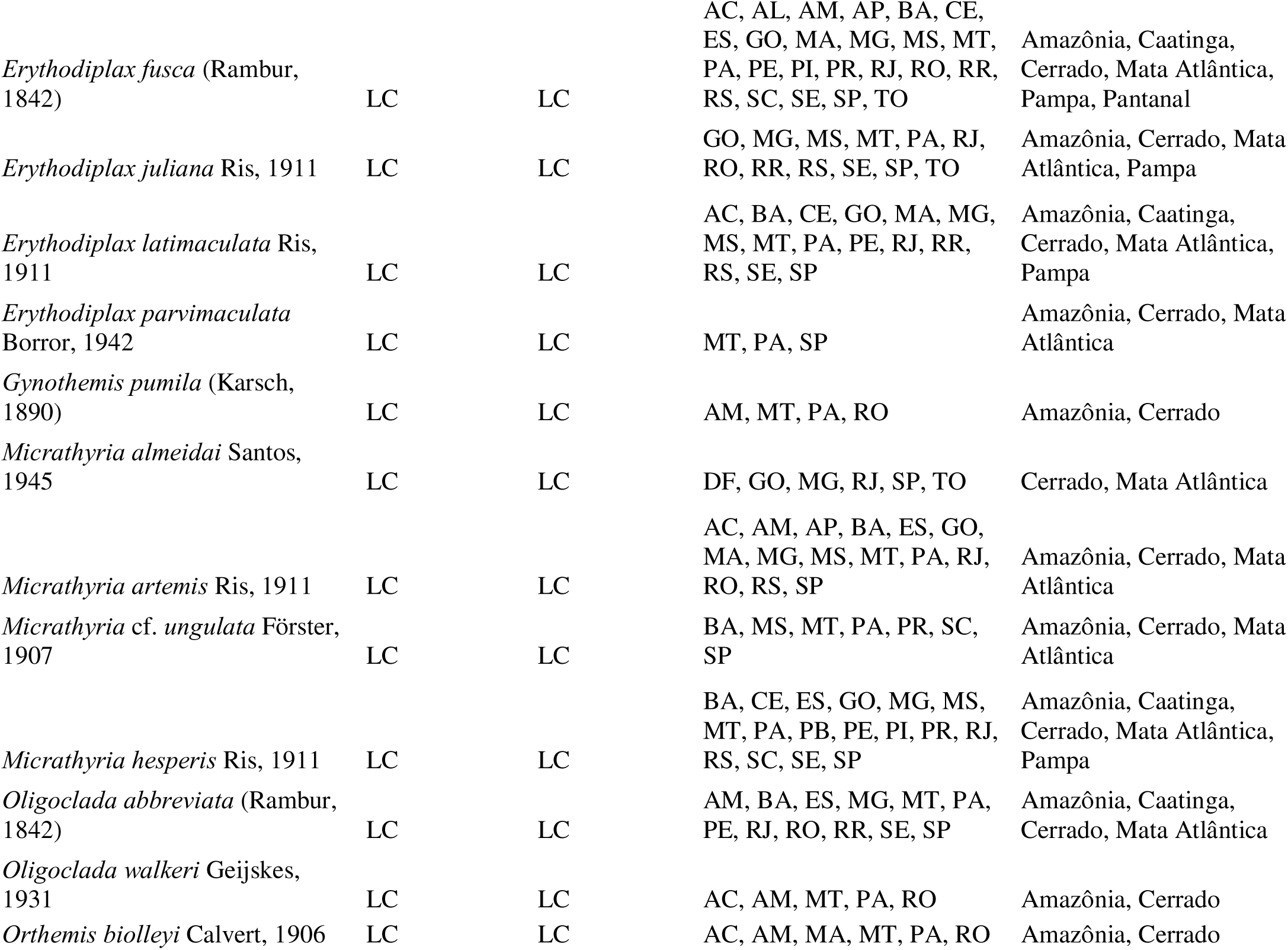
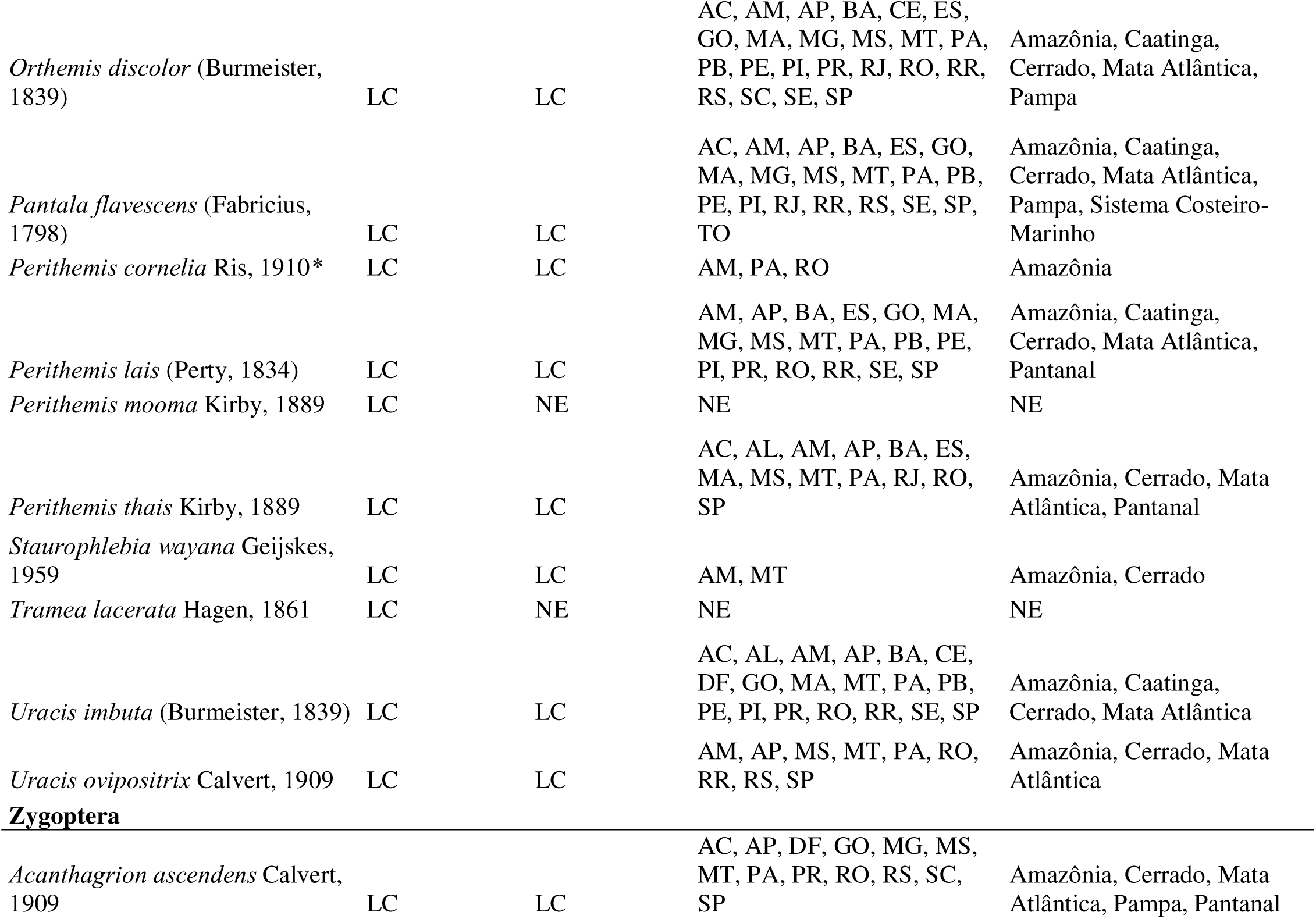
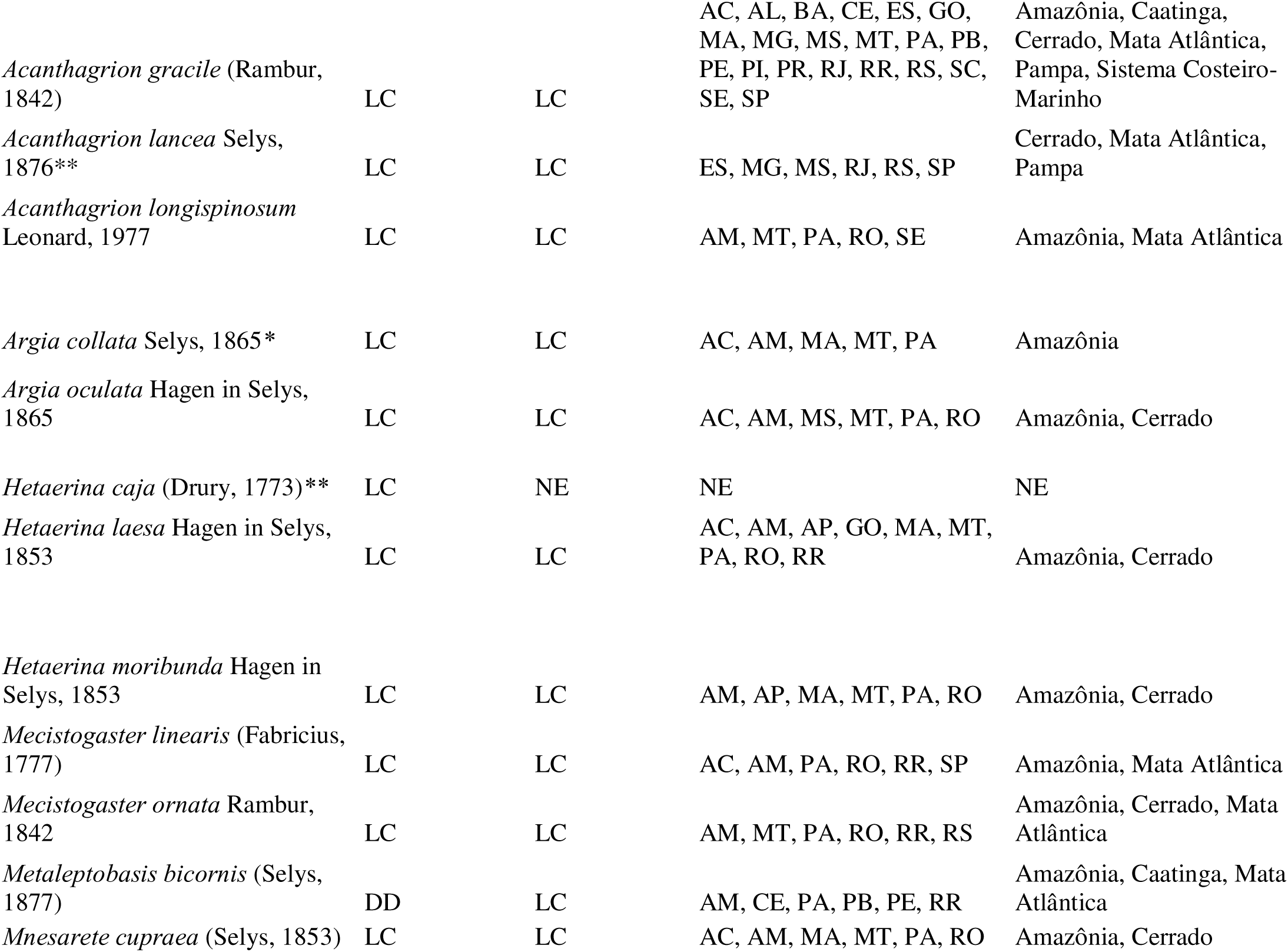
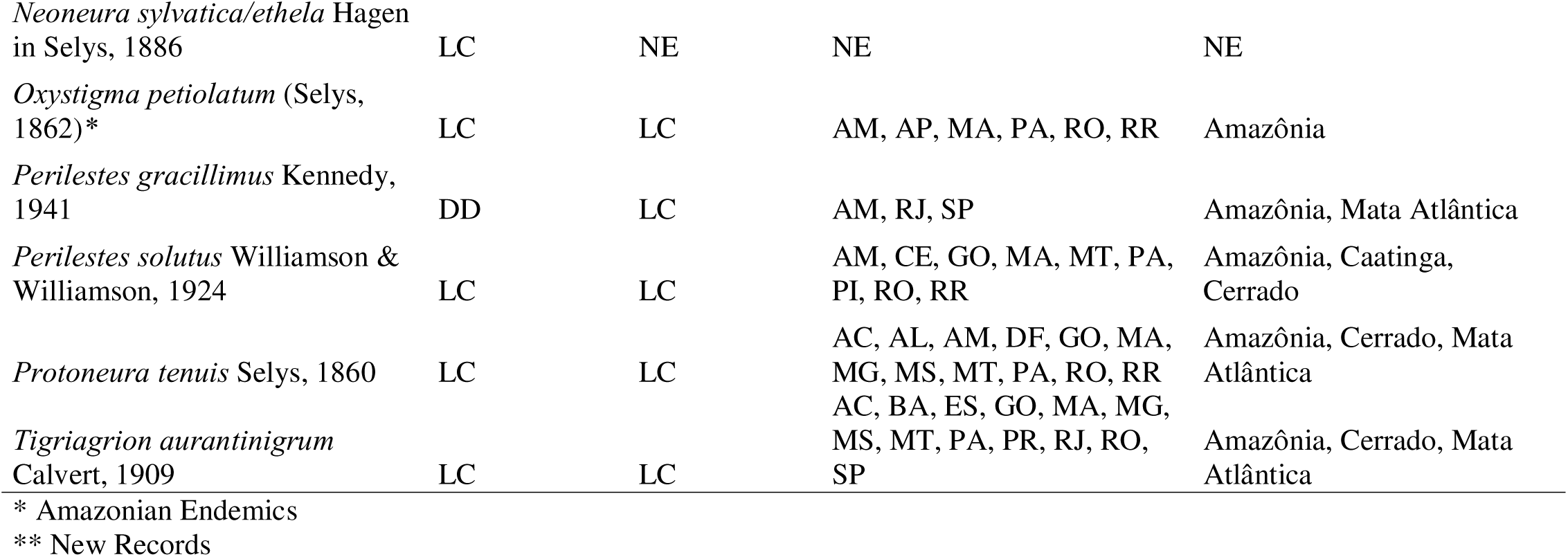
Species occurrence records by sampling site, including national distribution records and Red List threat status classifications. LC = low concern; NE = Not Evaluated. TI = Indigenous Territory; UF = Federative Units.

### Influence of environmental and landscape predictors on Odonata

The variance partitioning at the order level indicated that local variables explained a higher fraction of the total variance (Adjusted R² = 0.08), followed by landscape variables (Adjusted R² = 0.014). A large proportion of the variance remained unexplained (0.871), suggesting the influence of additional unmeasured factors. No spatial filter was retained by the forward selection. For suborder Zygoptera, the variance partitioning indicated that fine-scale spatial filters explained the higher fraction of the total variance (Adjusted R² = 0.084), followed by local variables (Adjusted R² = 0.046), and broad-scale spatial filters (Adjusted R² = 0.038). The unexplained variance was 0.836. Finally, for suborder Anisoptera, the variance partitioning showed that local and landscape variables explained approximately equal fractions of the total variance (Adjusted R² = 0.034; Adjusted R² = 0.030, respectively). The residuals were around 0.939.

The pRDA showed significant results only for order Odonata as a whole and for Zygoptera suborder. Anisoptera did not present any significant association with local habitat or landscape variables. For Odonata, we found a significant influence of local variables (AdjR² = 0.098; F = 1.654; p < 0,007). The first two ordination axes explained around 60.99% of the total variation (Fig. 2). Species such as *Mecistogaster* sp., *Hetaerina* sp., *Hetaerina laesa*, and *Hetaerina moribunda* showed a significant association with high values of HII and canopy cover (Fig. 2a). Besides, we found strong association of some *Erythrodiplax* species, *Acanthagrion gracile*, and *Hetaerina caja* with high values of macrophytes species richness, water depth and channel width of the streams (Fig. 2a).

**Fig. 2.**
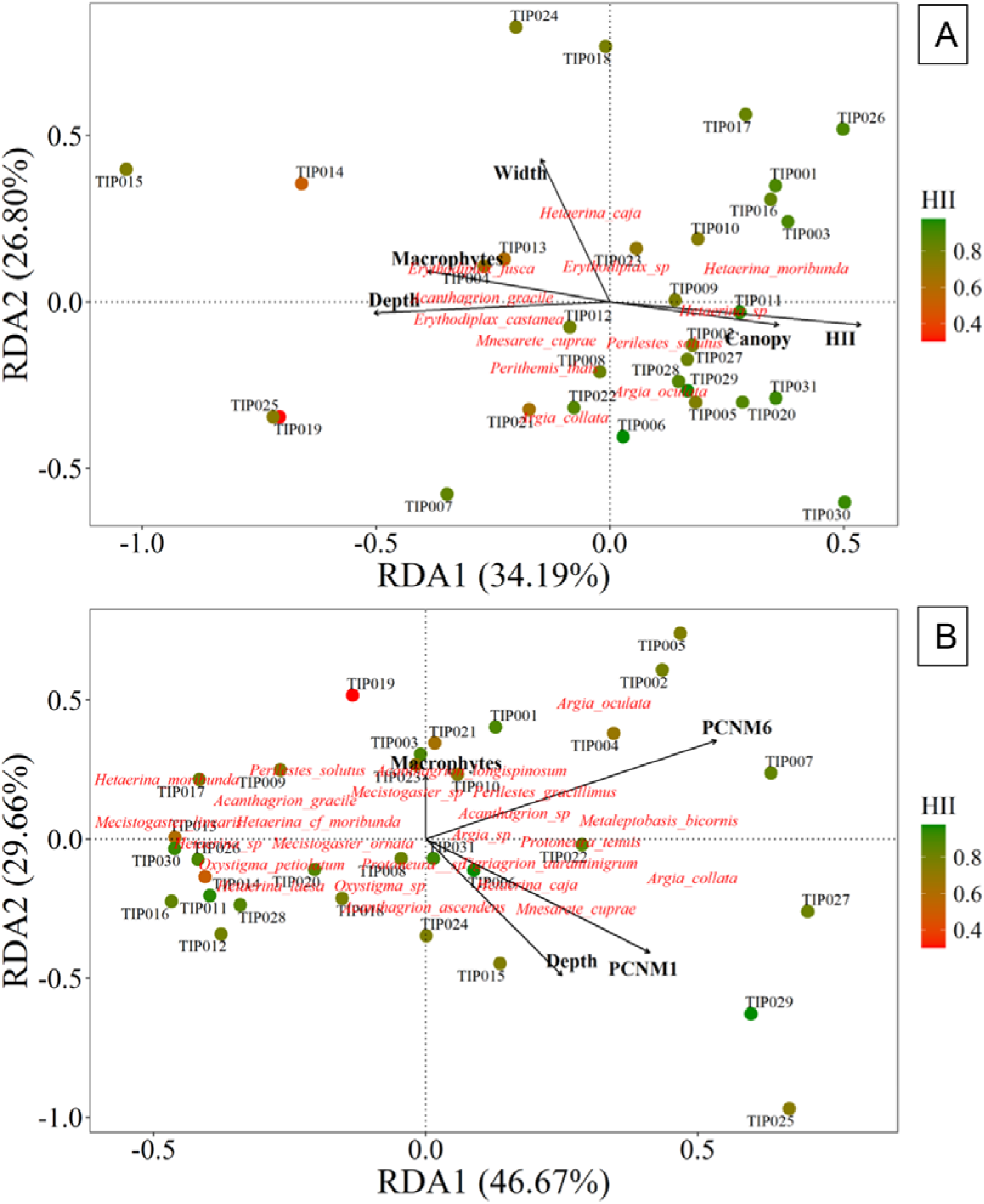
Triplot displays the relationship between local environmental variables and species composition of Odonata (order) and Zygoptera (suborder), with streams’ labels. Canopy = Local Forest (%); Depth and Width = physical structure of the streams’ channels (measured in centimeters and meters, respectively); HII = Habitat Integrity Index; Macrophytes = species number; PCNM - Principal Coordinates of Neighbour Matrices spatial filters. The color gradient ranges from reddish (lower habitat integrity) to greenish (higher habitat integrity).

Regarding Zygoptera responses, we found that local and broad- and fine-scale spatial predictors presented significant influence (AdjR² = 0.045; F = 1.786; p = 0.032; AdjR² = 0.115; F = 2.936; p < 0.001, respectively). The first two ordination axes explained around 76.33% of the total variation (Fig. 2). Species from genus *Acanthagrion*, *Argia*, *Metaleptobasis*, and *Perilestes* were associated with the fine-scale spatial filter (Fig. 2b). Additionally, species of *Argia*, *Tigriagrion aurantinigrum*, and Calopterygidae were associated with broad-scale spatial filters (Fig. 2b). Macrophyte species richness exerted a weaker influence on *Acanthagrion longispinosum* and *Mecistogaster* sp., whereas water depth exerted a stronger effect on species distribution patterns (Fig. 2b).

The TITAN analysis identified seven species of Odonata exhibiting significant association with environmental gradients (both local habitat and landscape) (Fig. 3). Species responses were detected along both landscape gradients (anthropic uses and forest) and local habitat gradients (number of macrophytes and canopy cover) (Fig. 3). Among Anisoptera, *Perithemis thais* showed a positive association with the anthropic gradient (Z+) and negative association with the forest gradient (Z−) (Fig. 3a). *Erythrodiplax fusca* exhibited a positive association with local forest (Z+) (Fig. 3d). For Zygoptera species, *Acanthagrion gracile* responded positively to macrophytes species richness (Z+). *Argia collata*, *Argia oculata* and *Hetaerina laesa* exhibited a negative association with forest gradient (Z−). *Perilestes solutus* presented positive association with local forest (Fig. 3d). All species-specific responses, including purity and reliability are summarized in Table S2.

**Fig. 3.**
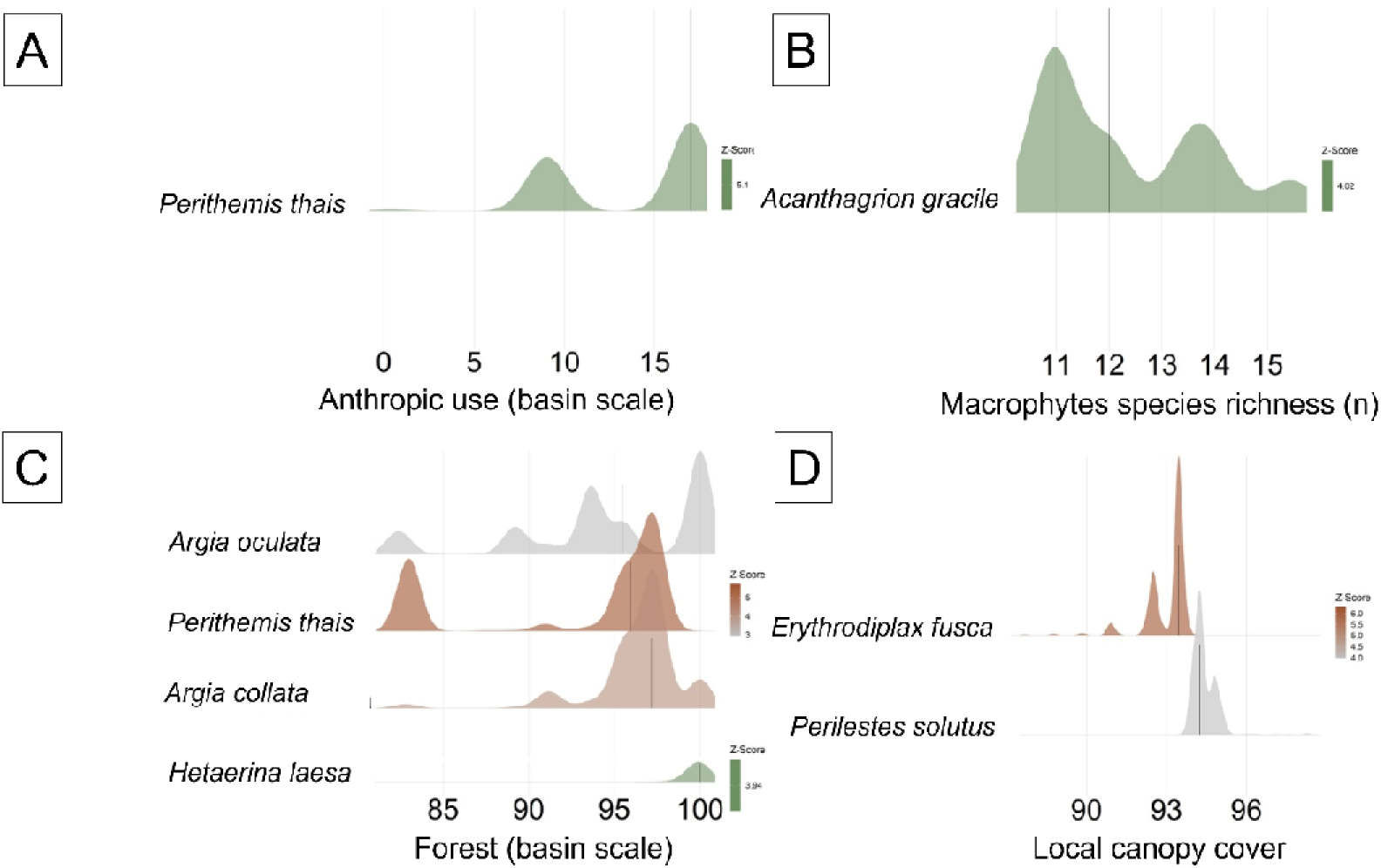
Ecological thresholds of Odonata species along environmental gradients, represented by local habitat and landscape components. Grey to green gradient indicates positive associations with environmental gradients (Z+); grey to brown gradient indicates negative associations with environmental gradients (Z−). A= Anthropic use; B= Macrophytes species richness; C= Forest; D= Local canopy cover.

## Discussion

Our hypotheses were partly corroborated, as significant relationships were detected for overall Odonata order and Zygoptera suborder, whereas Anisoptera did not show any association with the predictor’s dataset at the community level. These results provide clear evidence that even within well-preserved Indigenous Territories, Odonata assemblages are structured by a combination of local habitat filtering and spatial processes, with distinct mechanisms operating between suborders. This pattern suggests that the structuring mechanisms operating within the territory differ between suborders, reflecting their contrasting ecological and dispersal characteristics, as well as the integrity of streams within the Indigenous territory. The predominance of local habitat influence (macrophytes and other physical structures of the streams) on Odonata indicates an overall ecological pattern associated with riparian integrity. Moreover, the species-specific ecological thresholds revealed additional patterns that were not fully captured by community-level analyses. When both suborders were analyzed together, local environmental gradients emerged as the primary drivers of community structure; however, separating suborders revealed distinct ecological responses that would otherwise remain obscured.

Anisoptera and Zygoptera suborders differ substantially in ecological and physiological requirements (Oliveira-Junior et al. 2015; Miguel et al. 2017; O’Malley et al. 2020), and these distinctions are reflected in the roles of local and spatial components. Therefore, this can explain our results for both suborders, and the lack of significant outcomes for Anisoptera. This suborder is usually composed of species that are more tolerant of environmental modifications and present more effective dispersal movements compared to Zygoptera (May 1976; Brasil et al. 2017; Ferreira et al. 2023). Such ecological flexibility likely reduces the strength of detectable associations with environmental and spatial predictors at the community level, even though species-specific responses may still occur (Bastos et al. 2021; Brito et al. 2020a). Nevertheless, species-level threshold analyses demonstrated that certain Anisoptera taxa do respond to specific gradients, reinforcing the importance of complementary analytical approaches.

Considering Odonata and Zygoptera, there occurred significant prevalence of local habitat for the first, and spatial filters, for the last. The local habitat variables are associated with the provision of physical structure surrounding the stream channels, incidence of sunlight, and primary productivity (Oliveira-Junior et al. 2015; Calvão et al. 2023; Ferreira et al. 2024). Associations between species and environmental variables reinforce the influence of habitat structure and riparian integrity in shaping Odonata communities. Species such as *Hetaerina* sp., *H. caja, Mecistogaster* sp., *H. laesa*, and *H. moribunda* were consistently associated with higher values of habitat integrity and canopy cover, indicating strong dependence on well-preserved riparian environments. These results are consistent with previous studies showing that forest-specialist taxa are highly sensitive to canopy removal and habitat simplification (Díaz-Flórez et al. 2018).

Additionally, broad and fine scale spatial filters explained a higher fraction of Zygoptera variance, indicating dispersal constraints and source-sink dynamics. Some Zygoptera species (from genus *Mnesarete*, *Mecistogaster*, and some *Argia*) are highly dependent of local environmental conditions, and present less robust flying characteristics (e.g., narrow wings, long abdomens, which increases the inertia momentum, less robust thorax muscles), which prevents longer dispersal movements (Wootton 2020). These traits increase their sensitivity to spatial configuration and connectivity among habitats, reinforcing the role of spatial processes in structuring their assemblages. Besides, the higher vegetation cover and environmental stability in the indigenous land, provided by the riparian vegetation, might maintain some habitat connectivity, allowing species’ displacements across the landscapes (Leibold et al. 2004; Heino et al. 2015). This may explain the association between the species of the genus *Perilestes, Metaleptobasis*, and *Protoneura* and with the fine-scale spatial predictor.

Physical structures, such as bushes, small trees, wood debris, and leaf packs, are used by odonate males for territorialism, breeding, and foraging behaviors, and by females as oviposition sites (Resende et al. 2021). The mean local canopy cover was around 0.94, while the mean width was 2.4m, indicating shading conditions and a high presence of trees, which translate into more allochthonous resources and temperature stability, crucial conditions for more sensitive and forest specialists, such as Zygoptera. Such conditions enhance microclimatic stability, increase the availability of allochthonous resources, and create suitable microhabitats for sensitive species. *Hetaerina moribunda* exhibited wide distribution across the study area, with 150 individuals recorded in 26 of the 31 sampled sites. This broad occurrence suggests ecological plasticity within preserved forest environments, particularly under high canopy cover. Although *H. moribunda* is typically associated with Amazon and Cerrado biomes (ICMBio 2026), its frequency in structurally heterogeneous and well-preserved habitats reinforces its affinity for stable riparian conditions (Monteiro-Junior et al. 2014). However, community-level analyses alone did not fully capture the variability in species responses, underscoring the importance of threshold detection approaches (Silva et al. 2024; Sousa et al. 2025).

Additionally, *A. gracile* exhibited a positive ecological threshold for macrophyte species richness, becoming more frequent above specific levels of structural complexity. However, the peak abundance of this species occurred before the maximum macrophyte richness, indicating an intermediate response to the environmental gradient. *A. gracile* appears to benefit from increased structural complexity (macrophytes) under partial canopy opening, while higher macrophyte richness associated with severe canopy loss (Fares et al. 2020) leads to reduced abundance. These species-specific responses emphasize that ecological thresholds operate even in well-preserved Indigenous Territories and other protected areas and may serve as early indicators of environmental change (Brito et al. 2024a; Sousa et al. 2025).

In this context, ecological threshold analysis complemented multivariate results by identifying abrupt changes in species occurrence along environmental gradients. Anisoptera included two species with significant threshold responses (*E. fusca* and *P. thais*), whereas Zygoptera exhibited five responsive species (*A. gracile*, *A. collata*, *A. oculata*, *H. laesa*, and *P. solutus*). Although diversity patterns of both suborders in small tropical streams are well documented (Juen and De Marco 2011; Oliveira-Junior et al. 2015; Brasil et al. 2018; Brito et al. 2024b; Miguel et al. 2017), some species displayed patterns that differed slightly from general expectations.

Species of *Acanthagrion* typically occur across a broad range of environmental conditions (Fulan et al. 2011; Veras et al. 2022; Silva Junior et al. 2023), suggesting tolerance to microclimatic variation. This may explain the increased frequency of *A. gracile* along the macrophyte cover gradient. The highest occurrence of *A. gracile* was recorded at site TIP14, where the HII indicates moderate environmental alteration, potentially reflecting greater canopy openness and temperature variation (Oliveira-Junior et al. 2015). Additionally, aquatic macrophytes provide perching sites and refuge for larval stages (Fares et al. 2020), reinforcing the ecological relevance of structural heterogeneity.

The ecological threshold detected for *E. fusca*, characterized by increased frequency along the canopy cover gradient, may be associated with the presence of small canopy openings along streams. As a heliothermic species, *E. fusca* depends on solar radiation for thermoregulation (De Marco and Resende 2002; Vinagre et al. 2024). The highest abundance of *E. fusca* also occurred at TIP14, where greater sunlight incidence may favor population establishment. *Perithemis thais* showed a positive association with anthropic gradients and a negative association with forest cover, consistent with its preference for more open environments and direct solar radiation (De Marco and Resende 2002). This pattern indicates that even within predominantly conserved territories, local variation in landscape can influence species distribution.

Zygoptera displayed more consistent responses to environmental gradients, with significant associations to local habitat variables, particularly water depth and macrophyte richness. These findings reinforce the expectation that Zygoptera are more specialized and sensitive to local habitat conditions, whereas Anisoptera exhibits more generalist patterns detectable primarily at the species level. Such differentiation highlights the importance of analyzing suborders separately when investigating freshwater insect assemblages.

The environmental characteristics of the streams within the Panará Indigenous Territory indicate high overall habitat integrity, as evidenced by elevated forest cover and HII values. However, variation in macrophyte numbers, water depth, and surrounding agricultural land use demonstrates meaningful environmental heterogeneity among sites. This internal heterogeneity is likely a key factor sustaining high biodiversity, as it provides a mosaic of microhabitats and ecological conditions that support species with different ecological requirements.

The relatively high proportion of unexplained variance observed in our models is consistent with other ecological studies in tropical freshwater systems and likely reflects the influence of additional factors not included in our analyses, such as water physicochemical properties, temporal dynamics, biotic interactions, and microhabitat complexity. This highlights the inherent complexity of Amazonian stream ecosystems and suggests that multiple interacting processes operate simultaneously to shape biodiversity patterns.

Our records indicate a superpopulation of specimens identified as H. caja, whose taxonomic status, according to ICMBio, is recorded only for the states of Amazonas and Roraima. Similarly, *A. lancea*, which has only been recorded for the Cerrado, Atlantic Forest, and Pampa biomes, was recorded for the first time on the Amazon. Also, three endemic Amazonian species were recorded: *P. cornelia*, A. *collata*, and *O. petiolatum*, reinforcing the importance of preserving the territory for the protection of these species.

Beyond the ecological patterns detected, our findings reinforce the urgent need to strengthen collaborative conservation strategies that actively integrate Indigenous Peoples as central actors in biodiversity research and management (Sze et al. 2022; Osborne et al. 2024). The conservation of Amazonian freshwater ecosystems increasingly depends on the combined efforts of academic institutions, conservation agencies, and Indigenous communities, whose territories harbor some of the most well-preserved aquatic systems in the region. Indigenous Peoples possess detailed ecological knowledge derived from long-term interactions with their environments, which, when combined with scientific approaches, enhances the detection of subtle environmental changes, improves monitoring effectiveness, and supports more resilient conservation outcomes. In a context of accelerating environmental change, biodiversity conservation can no longer rely on isolated efforts; rather, it requires genuine partnerships that recognize Indigenous Peoples not only as guardians of biodiversity, but as co-producers of knowledge and key allies in sustaining ecological integrity across Amazonian landscapes.

## Conclusions

In summary, our findings indicate that Odonata assemblages in the Panará Indigenous Territory are shaped by both habitat filtering and spatial processes, whose relative importance varies between suborders. Local habitat integrity and stream structure were more important for overall Odonata, whereas Zygoptera showed stronger spatial structuring, likely reflecting dispersal constraints and higher environmental specialization. In contrast, Anisoptera responses were weak at the community level but detectable for species, emphasizing that assemblage-wide analyses may mask species-specific environmental thresholds. These findings reinforce that Indigenous Territories not only preserve habitat integrity but also maintain ecological processes and environmental heterogeneity necessary to sustain biodiversity patterns and species-specific responses. By integrating multiscale environmental predictors with species-specific threshold responses, this study advances the understanding of how biodiversity is structured in highly conserved Amazonian landscapes, providing a framework for biomonitoring and conservation planning.

## Supporting information

Tabela Suplementar 1

## Acknowledgments

We also acknowledge the support of the projects INCT Sínteses da Biodiversidade Amazônica (Process No. 406767/2022–0), Programa de Pesquisa em Biodiversidade da Amazônia Oriental – PPBio AmOr (Process No. 441257/2023–2), and Programa Ecológico de Longa Duração da Amazônia Oriental PELD-AmOr (Process No. 445970/2024–3). We thank the field team from the Federal University of Pará and the Conservation International team for their logistical and technical support during the field expeditions conducted in the Panará Indigenous Territory. We extend our special gratitude to the Panará Indigenous researchers, whose active participation throughout all stages of the study, from methodological discussions and selection of sampling sites to fieldwork, specimen triage, and the sharing of ecological knowledge, was fundamental to the development and quality of this research. We are grateful to the Universidade Federal do Pará (UFPA) for providing physical infrastructure and institutional support for specimen identification and data analysis. This manuscript is the result of the doctoral research of the first author (JCMF), a CAPES scholarship holder (Process No. 88887.176346/2025-00), and was developed within the context of the Scientific Article Workshop promoted by the Laboratório de Ecologia e Conservação (LABECO) in conjunction with the Programa de Pós-graduação em Ecologia (PPGECO). We acknowledge CNPq for postdoctoral fellowship to JSB (Process No. 151038/2024-4), and GMC (process 88887.939579/2024-00). Finally, we thank the participants of the 1st Workshop for Discussion and Writing of Basic Biology Articles on Aquatic Insects, held at the Federal University of Pará (UFPA) in January 2026, for valuable discussions and contributions.

## Author contributions

All authors conceptualized the study. JCMF and JSB wrote the first draft of the manuscript. LFAM, LJ and TSM sourced funding. KKP, KkP, SeP, SaP, KwP, KeP, SpP, NP, PPP, PP, TP collected and prepared the material. AROC, BC IMAE, LS, PJ and RP support and logistics in the field. EP, JVAS, PGSG and YCCD collected, prepared the material and data analyses. BLS, GMC, FFB, FMBS, FSS and RL analyzed the data. All authors reviewed and agreed with the final version of the manuscript.

## Declarations

## Consent to participate

The authors have consented to the submission.

## Ethics approval

The capture, collection, and transport of biological material were authorized by the Instituto Brasileiro do Meio Ambiente e dos Recursos Naturais Renováveis (IBAMA; SISBIO permit No. 4681-1) and approved by the Animal Use Ethics Committee of the Federal University of Pará (CEUA No. 8293020418). The collection of the biological samples was authorized by the Biodiversity Authorization and Information System (SISBIO) of the federal Chico Mendes Institute for Biodiversity Conservation (ICMBio), through permanent license 11841-2. Authorization to access Indigenous territories was granted by the Fundação Nacional dos Povos Indígenas (FUNAI) under administrative process No. 08620.012263/2023-71.

